# Role of microstructure of cellulosic mucilage in seed anchorage: A mechanical interpretation

**DOI:** 10.1101/2024.01.08.574694

**Authors:** Krithika Bhaskaran, Puchalapalli Saveri, Abhijit P. Deshpande, Susy Varughese

## Abstract

Cellulosic, hemicellulosic and pectinaceous mucilages produced by certain angiosperms as adaptation in *myxodiaspory* are investigated in the past for seed dispersal. The present understanding of *zoochory* and *telechory* are based on mucilage amount, state of hydration and to a limited extent, role of mucilage microstructure studied using adhesion and friction. Pectinaceous mucilages have less adhesion and supports dispersal by *zoochory*. However, in the case of cellulosic mucilages, the role played by the cellulosic fibrils in seed dispersal is not clear, especially, since they have a negative correlation with *endozoochory*. Using fresh cellulosic seed mucilages from, sweet basil (*Ocimum basilicum*) and chia (*Salvia hispanica*) we investigate the role of microstructure of the mucilage in two key behaviours: anchoring and adhesion properties of the seeds through rheology. We report a special large deformation mechanism triggered through ‘strain stiffening’ operational in these cellulosic mucilages. In many biopolymers semi-flexible polymer chains and other aligning elements contribute to the strain stiffening. However, the strain stiffening and strong wet adhesion observed in these mucilages have a significant role from the cellulosic components. This behaviour is more pronounced in basil seeds and presents a plausible structure-property mechanism for *anti-telechory* favoured by plant species found in arid habitats.

## 1 Introduction

Plants adapt various unique strategies for their survival in diverse environments. Myxodiaspory is one of those strategies shown by seeds/achenes of angiosperms [1]. Myxodiaspory refers to the exudation of mucilage, a gelatinous complex polysaccharide upon imbibition of water by the seeds. The polysaccharides present in the seed/achene mucilage are constituted of pectin, cellulose, hemicellulose and starch, with varying combinations and compositions across different species to suit the seed dispersal mechanism [2–4]. The mucilage is a highly functional trait in seed/achene physiology which helps in seed dispersal, hydration, anchorage, germination and also in early seedling development [3, 5, 6]. It also helps in protecting the structural integrity of the seed by acting as a protective sheath surrounding the seed [7] in addition to regulating the seed germination [3, 6].

Different seed dispersal mechanisms such as anti-telechory, endozoochory and epizoochory are governed by the adhesive, frictional and mechanical properties of the mucilage [8–10]. Studies on endozoochoric dispersal of different mucilagenous seeds reveal that, cellulosic mucilages such as, *Ocimum Bacillicum* (sweet basil) and *Salvia hispanica* (chia) pass through the digestive system of pigeons but did not germinate whereas other hemicellulosic and pectinaceous mucilages such as, *Plantago Ovata* (plantago) and *Linum usitatissimum* (flax) facilitates the endozoochoric dispersal [8] by showing germination. Interestingly, mucilage secretion is shown to be selective response to the dislodgement by erosive forces during surface flows especially in cellulosic mucilages in semi-arid regions (*anti-telechory*) [10, 11]. Seed dislodgement studies on 52 different species under high surface flows with velocities ranging from 6.7 x 10^−5^ to 1.4 x 10^−2^ l/s cm [10] were carried out to establish that mucilage helps in anchoring the seed to substrate. This study has shown that higher mucilage mass and drying of mucilage helps in increased seed anchorage [10]. However, this study did not explore the role of mucilage micro structure in seed anchorage. Limited studies were reported on the role of mucilage micro structure in seed anchorage and dispersal mechanisms [8, 9, 12]. Adhesion studies conducted on hemicellulosic plantago and pectinaceous flax seed mucilage showed better adhesion for flax seed mucilage on glass substrate under wet and dry conditions [9, 12]. It is shown that the presence of hemicellulose in seed mucilage weakens the adhesion strength and frictional properties but still facilitates the zoochoric dispersal and seed anchorage [12]. Another study on cellulosic *Helianthemum violaceum* and pectinaceous *Fumana ericifolia* mucilages has shown that the presence of cellulose in seed mucilage increases the adhesion strength [13, 14] on glass. It is hypothesised that cellulose binds to the pectin in the mucilage providing additional strength to the mucilage and thus enhancing the anchorage to the soil. This suggested that cellulosic mucilage facilitates anti-telechory where the seed is anchored to the soil/substrate and restricts the long range dispersal by water run off. All these studies clearly indicates that the cellulose in mucilage microstructure has a certain role to play in its functioning. However, the exact contributions from the cellulosic fibrils to the mucilage response under shear forces remain unclear and needs to be elucidated under mechanical deformation conditions. In this context, a quantitative analysis of the structure-function relationship of the mucilage can help to understand the role of different components of the mucilage and their hierarchical structural organization in its mechanical response [15, 16].

Rheology is a widely used approach to investigate various material systems to understand their structural response to mechanical deformation (linear and non-linear shear) [17–20]. When subjected to large strains/deformations, one of the key characteristic response of certain soft biological materials is ’strain stiffening’ [21]. Some of the classic examples of such biological materials exhibiting strain stiffening include F-actin [22–24], fibrin [25–27], collagen [20, 27, 28], neurofilaments [29] which are of animal origin and alginate [19, 30], agarose [31] and pectin gels [17, 32] of plant origin. The strain stiffening response of these systems is also associated with their physiological functions and originates from the macromolecular network structure. Alginate and low-methoxy pectin get ionically crosslinked in the presence of divalent cations such as calcium (Ca^2+^) resulting in hydrogel network by forming egg-box dimers and bundles [17, 33, 34]. Low-methoxy pectin, which is also a major constituent of seed mucilages, is shown to exhibit strain stiffening behaviour under large amplitude oscillatory shear experiments. The strain stiffening behaviour of the pectin gels originates from the calcium-crosslinked polygalacturonan chains which form rigid rod-like egg-box bundles connected through semi-flexible polymer chains [17]. However, seed mucilages being a complex systems with not only pectin based networks, but also cellulose and hemicellulose components, may have different roles in their overall mechanical response. In this context, using rheological response of the mucilage as a tool, the following questions are addressed: 1) Does seed mucilage being also rich in pectin show strain stiffening response under mechanical deformations? 2) Does the presence of cellulose change the strain stiffening behaviour, if it is present? 3)What ecological functions are served by this response and the microstructure? 4) Does the cellulosic microstructure play any special role in telechory and seed anchorage to the soil?

In earlier studies, seed mucilages reconstituted from dried powder were used for characterizing the rheological behaviour in the context of food applications [35–39]. However, in those studies, the structure of the native mucilage was altered by one or more of the processes followed, such as, use of other chemicals, centrifugation and freeze-drying [40]. In the present study, to understand the *in vitro* behaviour of seed mucilage in plant ecology, fresh native mucilage from two widely studied myxodiasporic model species, sweet basil (*Ocimum basilicum*) and chia (*Salvia hispanica*) are investigated using rheology along with low-methoxy pectin gels for comparison purpose. Both small and large amplitude oscillatory shear (SAOS and LAOS) experiments and wet tack behaviour are specifically investigated to understand seed anchorage and telechory in these species.

## 2 Materials and methods

Sweet basil (*Ocimum basilicum*) and chia (*Salvia hispanica*) seeds were purchased from Essence Organics (Aurangabad, Maharashtra, India.) and Anveshan Farm Technologies Pvt. Ltd. (Gurugram, Haryana, India.) respectively. The seeds were certified organic to avoid chemical contamination. Low methoxy pectin (LMP) and high methoxy pectin (HMP) of citrus origin with a galacturonic acid content of 87% and 81% and degree of esterification 39% and 67% respectively were purchased from Krishna Pectins, Jalgaon, Maharashtra, India. Calcium chloride dihydrate (*CaCl*_2_.2*H*_2_*O*) used for cross-linking low methoxy pectin was procured from Merck Specialities Private Limited, Mumbai, Maharashtra, India.

### Collection of basil and chia seed mucilage

For collecting the mucilage, seeds were soaked in double distilled water at a seed-to-water ratio of 1:40 for 30 min at room temperature (25-30 °C). Seeds upon uptake of water secrete the mucilage. Excess water was filtered off using a nylon mesh and the swollen seeds were squeezed gently between two meshes to separate the mucilage from the seed surface. The separated mucilage was collected from the mesh surface. The collected mucilage was kept for degassing in a desiccator for 15 min to remove entrapped air. The mucilage was allowed to rest for 30 min before the rheological studies.

### Determination of solid fraction

The solid fraction of the mucilage gives an estimate of the total polysaccharide present in it. The solid fraction of the freshly collected seed mucilage was determined by drying the mucilage. Mucilage was spread on glass slides and placed in a vacuum oven at 27 °C and at a vacuum pressure of 50 mbar. The sample was weighed every hour until the weight remained constant. The solid content of the mucilage was calculated using the following equation,

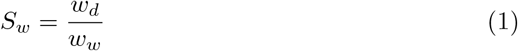

where *S*_*w*_ is the solid fraction in the mucilage, *w*_*w*_ and *w*_*d*_ are the initial (as collected) and final (as dried) weight of the mucilage. The polysaccharide content in freshly collected basil and chia seed mucilage is 0.8 wt% and 0.35 wt% respectively.

### Staining of swollen seeds

To ascertain the presence and location of pectin-rich and cellulose-rich regions in the seed mucilages, the swollen seeds were stained using Ruthenium red and Methylene blue dyes respectively. Simultaneous use of both the dyes in a sequence of ruthenium red followed by methylene blue gave the required preferential staining.

### Preparation of pectin gels

Pectin gels with different amounts of pectin and crosslinker concentrations were prepared for comparison with the mucilage. Low methoxy pectin gels were prepared according to the procedure reported in the literature [17]. Low methoxy pectin solutions of the required concentration were prepared in de-ionised water and stirred for 16-18 h at 60 °C for complete dissolution. Calcium chloride (*CaCl*_2_.2*H*_2_*O*) is used as a crosslinking agent. The stoichiometric ratio (R) relates the concentration of calcium ions to the non-methoxylated galacturonic acid content in the pectin. R is expressed

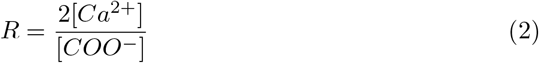

The extent of crosslinking was varied by varying the R value. Pectin-Ca gels were prepared for different R values by adding the prepared calcium chloride solution to the pectin solution drop by drop with stirring at 25-30 °C. The mixture was stirred for 10 min and left undisturbed for 24 h at room temperature (25-30 °C) for gel formation.

### Mechanical deformation behaviour of mucilage

Rheological investigations were carried out on freshly collected seed mucilages as well as on whole seeds swollen in water. These investigations are used to establish the gel-like response of the seed mucilages, adhesion and tack behaviour as well as the microstructural contributions to the observed mechanical responses. Rheological studies not only help in understanding the viscoelastic response of materials under mechanical shearing, it also sheds light on the microstructural contributions to the response. The storage modulus (G^*′*^(*ω*)) and the loss modulus (G^*′′*^(*ω*)) (hereafter referred to as G^*′*^ and G^*′′*^ respectively) describe the elastic and viscous response of the material and can be obtained from the oscillatory shear experiments carried out on the mucilage. Rheological studies were carried out on a stress-controlled rheometer, Anton Paar Physica MCR 301. A cone and plate geometry (CP 50-2/TG-SN5712) with a diameter of 50 mm, a true gap of 47 *µ*m and a cone angle of 1.995° was used. Oscillatory shear experiments were carried out to study both the linear and non-linear viscoelastic response of the mucilage. To study the linear deformation behaviour, small amplitude oscillatory shear (SAOS) experiments were performed at a strain amplitude (*γ*_0_) of 1% and angular frequency (*ω*) ranging from 0.1 to 100 rad/s. Large amplitude oscillatory shear (LAOS) experiments were carried out in the strain amplitude range 0.01 to 1000% at a constant angular frequency of 1 rad/s to study the non-linear deformation behaviour. All the measurements were carried out at a temperature of 25^°^C unless otherwise specified and were repeated thrice on separate mucilage samples for reproducibility. A silicone oil solvent trap was used to prevent the evaporation of water from the sample during the experiments.

### Analysis of the viscoleastic response

In the non-linear regime during a LAOS measurement, the stress response becomes a function of higher harmonics of the sinusoidal strain input. The overall moduli obtained from strain amplitude sweep contains the first harmonic information without reflecting the influence of higher harmonics. Analysis of intra-cycle stress-strain raw data using Lissajous-Bowditch (L-B) plots helps to understand the influence of higher harmonics on the material response [41, 42]. One approach for this analysis is based on strain stiffening index(S) that can be calculated from modulus at minimum and maximum strain values [41], while another approach known as Sequence of Physical Process (SPP) is based on calculation of transient moduli at every point on the L-B plots [43].

Non-linear behavior observed in L-B plots can be quantified by defining a parameter called strain stiffening index S, as a function of the minimum-strain elastic modulus 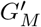 and the maximum strain elastic modulus 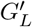 using the equation

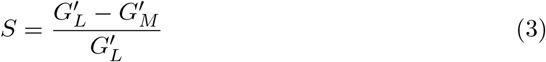

where, 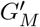 - tangent modulus at minimum strain (*γ* = 0) and 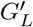 - secant modulus at maximum strain (*γ* = *γ*_0_) during one cycle at a given strain amplitude (Figure 1(a)). S = 0 implies linear response, where the material functions do not depend on the strain. 0 *<* S *<* 1 indicates strain stiffening and S *<* 0 indicates strain softening behaviour of the material system [41]. Sequence of Physical Process (SPP) analysis estimates transient moduli from the L-B plots by calculating the slope of the L-B curve at each point in the cycle for a given strain amplitude. The transient or instantaneous elastic modulus 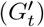 is obtained from the slope of stress-strain curve in L-B plot and the slope values obtained from stress-strain rate curve in L-B plot implies transient loss modulus 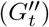. The transient moduli is defined as,

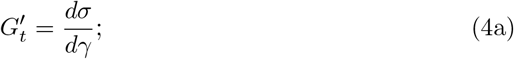

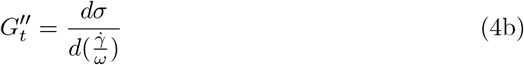

**Fig. 1.**
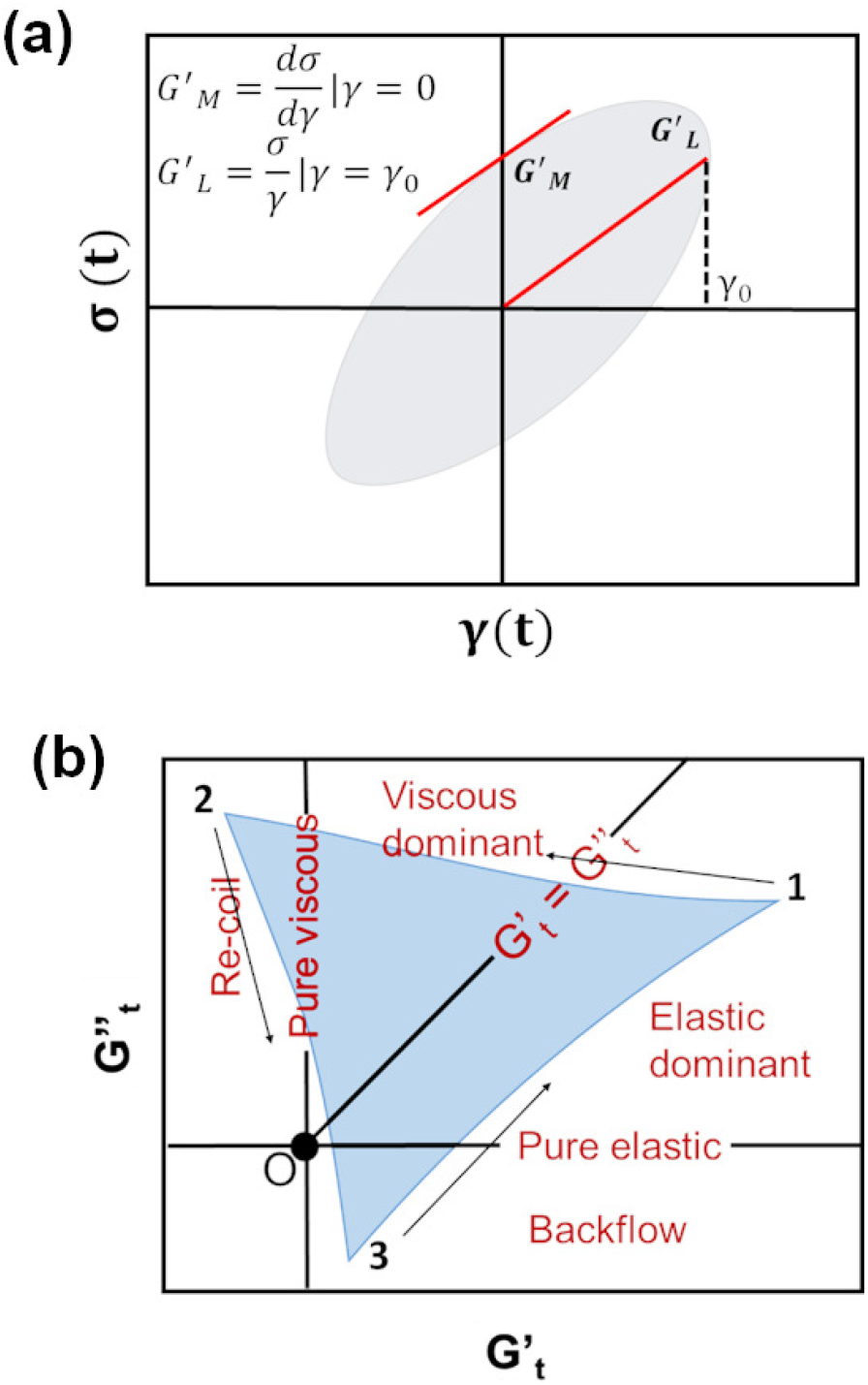
(a) Model Lissajous-Bowditch (L-B) plot - stress-strain cycle data with tangent and secant moduli at minimum and maximum strain [41] (b) SPP analysis - Position of different regions in the Cole-Cole plot of transient moduli[43]

The earlier mentioned moduli 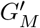 and 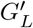 imply the response only at limiting conditions of minimum (*γ* = 0) and maximum strain (*γ* = *γ*_0_), whereas, the transient moduli encompasses the deformation response continuously along the L-B curve. This allows to obtain detailed transient information related to non-linear features such as, strain stiffening and strain softening [44].

The Cole-Cole plot of the estimated transient moduli 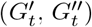 is analysed by classifying it into different regions as shown in Figure 1(b). The horizontal line and vertical line passing through the origin indicate purely elastic and purely viscous behaviour respectively. The 45° line through the origin represents the flow condition where the transient storage modulus 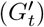 is equal to the transient loss modulus 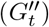. The positive region above and below the flow condition represents viscous and elastic dominant regions respectively. The region between positive 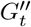 and negative 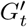 represents viscous recoil and the region between negative 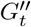 and positive 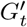 represents backflow [43–45].

### Tack response of seed mucilage

Tack experiments on swollen seeds and pectin gels were performed on the rheometer using parallel plate of 25 mm dia with roughened surface. On the lower platen of the rheometer, a single layer of swollen basil (47 seeds) and chia seeds (53 seeds) with mucilage envelope were loaded at 3 mm and 2.5 mm gap respectively. The gap was chosen based on the size of the swollen seeds so that the top plate comes in contact with the gel on the seed surface without force application. Similarly, basil sed mucilage and chia seed mucilage equivalent pectin gels (pectin gel 0.5 wt%, R=0.5 and pectin gel 0.35 wt%, R=0.5) were also loaded at 3 mm gap. Then, the tack experiment was performed by lifting the top plate at a retraction speed of 10 *µ*m s^−1^. The negative normal force (mN) exerted by the swollen seeds and pectin gels during retraction was measured. The adhesive strength was obtained from the normal force vs. time (s) plot.

## 3 Results

### 3.1 Viscoelasticity of seed mucilages - weak hydrogels?

The linear rheological analysis using the variation of storage (*G*^*′*^(*ω*)) and loss modulus (*G*^*′′*^(*ω*)) with angular frequency (*ω*) is used to understand the physical nature of the gel. Small amplitude oscillatory shear (SAOS) response of freshly collected basil seed mucilage (BSM) and chia seed mucilage (CSM) is shown in Figure 2.

**Fig. 2.**
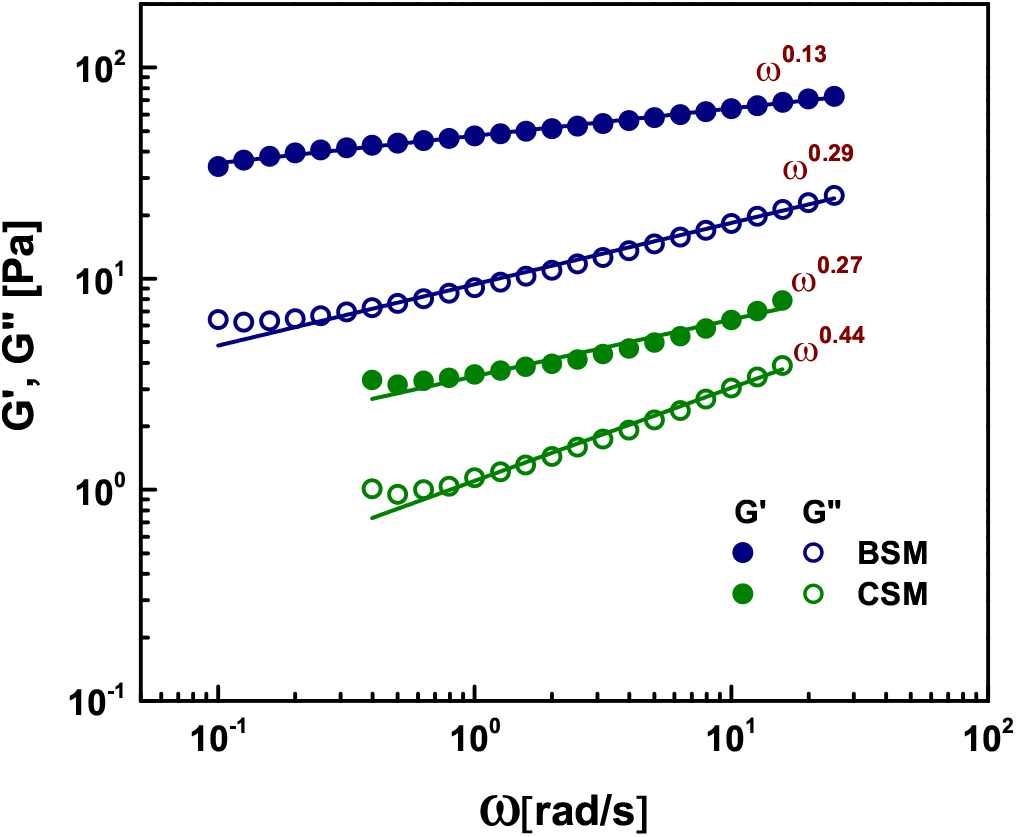
Small amplitude oscillatory shear response of BSM and CSM at 1% strain amplitude

For both the mucilages, the storage modulus is higher than the loss modulus throughout the frequency range studied indicating viscoelastic solid-like behaviour. BSM shows higher storage and loss moduli than CSM at all the frequencies. The storage and loss moduli of both the mucilages show a frequency dependence, *G*^*′*^ ∝ (*ω*)^*a*^ and *G*^*′′*^ ∝ (*ω*)^*b*^. A power law fitting of the data gives *a* and *b* in the range of 0.1 - 0.4 indicating the weak frequency dependent nature of moduli exhibited by weak gels [46]. In comparison to reconstituted mucilages from dry powders, freshly collected mucilages have lower storage modulus. This could be due to the higher solid content in the reconstituted mucilage and also the structural changes due to processing [39]. The small amplitude oscillatory shear (SAOS) analysis is limited to studying the general nature of these biopolymers since the response is linear in this range. To understand the contributions of microstructure and the structural changes in the mucilage during the mechanical deformation, large amplitude oscillatory shear (LAOS) experiments were carried out. The variation of storage (*G*^*′*^(*γ*_0_)) and loss (*G*^*′′*^(*γ*_0_)) moduli with strain amplitude for both basil and chia mucilages is shown in Figure 3. The overall response can be classified into four different strain regions. Region I: *γ*_0_ ≦ 2%, linear response region where both *G*^*′*^ and *G*^*′′*^ are independent of strain. Region II: 2% *< γ*_0_ ≦ 40%, the region where *G*^*′*^ tends to decrease with increasing strain amplitude indicating non-linear response. Region III: 40% *< γ*_0_ ≦ 200%, the cross-over region where (*G*^*′*^) and (*G*^*′′*^) crossover. For BSM, this occurs over *γ*_0_ of 100-125 % and for CSM, it is at 150%. Region IV: transition to viscoelastic fluid-like behaviour occurs beyond the crossover at *γ*_0_ *>* 200%, (where *G*^*′′*^ *>* G^*′*^).

**Fig. 3.**
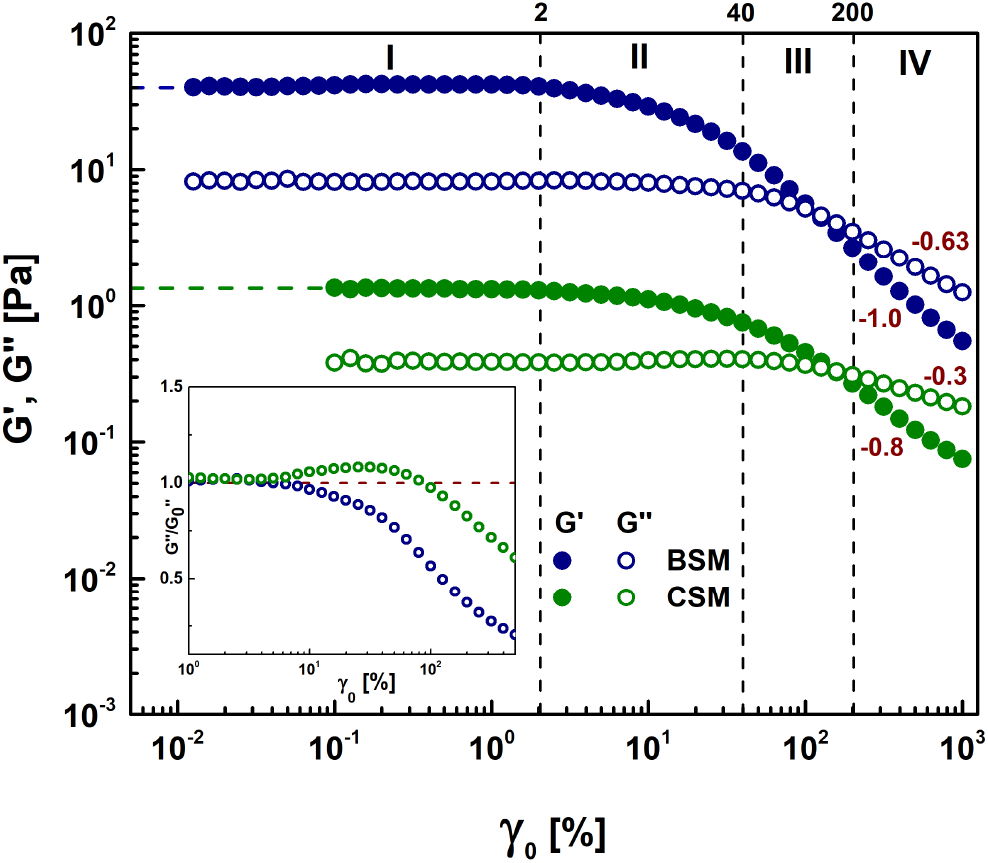
LAOS response of BSM and CSM at angular frequency of 1 rad/s. The vertical lines indicate the strain amplitude limits for the four rheological regions. The horizontal lines indicate the values of the mucilages. The inset in the figure 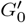 shows the overshoot in G^*′′*^ observed for chia seed mucilage.

The zero shear modulus 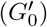 for BSM is one order of magnitude higher than that of CSM during LAOS. BSM shows no overshoot in *G*^*′′*^ while a weak overshoot is observed for CSM as shown in the inset in Figure 3. BSM with no overshoot represents type I material system in which polymer chains/microstructural entities align in the direction of flow [47]. On the other hand, CSM with weak overshoot represents type III materials with additional rearrangement and formation of temporal networks in the microstructure during deformation [48].

### 3.2 How close are seed mucilages to pectin gels?

#### 3.2.1 Structural composition

The FT-Raman spectra for BSM and CSM along with pectin gels and low-methoxy pectin are shown in Figure 4. The peaks at 2950 (*ν*(C-H)), 1750 (*ν*(C=O), ester), 1459(C-H_2_) and 1098 (assymmetric glycosidic bonds) cm^−1^, which corresponds to pectin, can be observed in both the seed mucilages as well as in the low-methoxy pectin and pectin gels [49]. The presence of crosslinked pectin in both basil and chia seed mucilages is well established in the literature [50–52]. The slight shift in these peaks are expected due to the differences in the composition of the pectins. In addition, both the seed mucilages exhibit distinct peaks at 1378 (*ω*(C-H_2_)), 897 (glycosidic bonds) and 560 (*β*(O-C-O)) cm^−1^ which shows the presence of hemicellulose and cellulose [53]. Basil seed mucilage, especially shows peaks at 438 (*ν*(CCO)ring) and 361 cm^−1^, that are characteristic of cellulose presence [54]. FT Raman analysis confirms that seed mucilages have crosslinked pectin along with hemicellulose and cellulose in their microstructure.

**Fig. 4.**
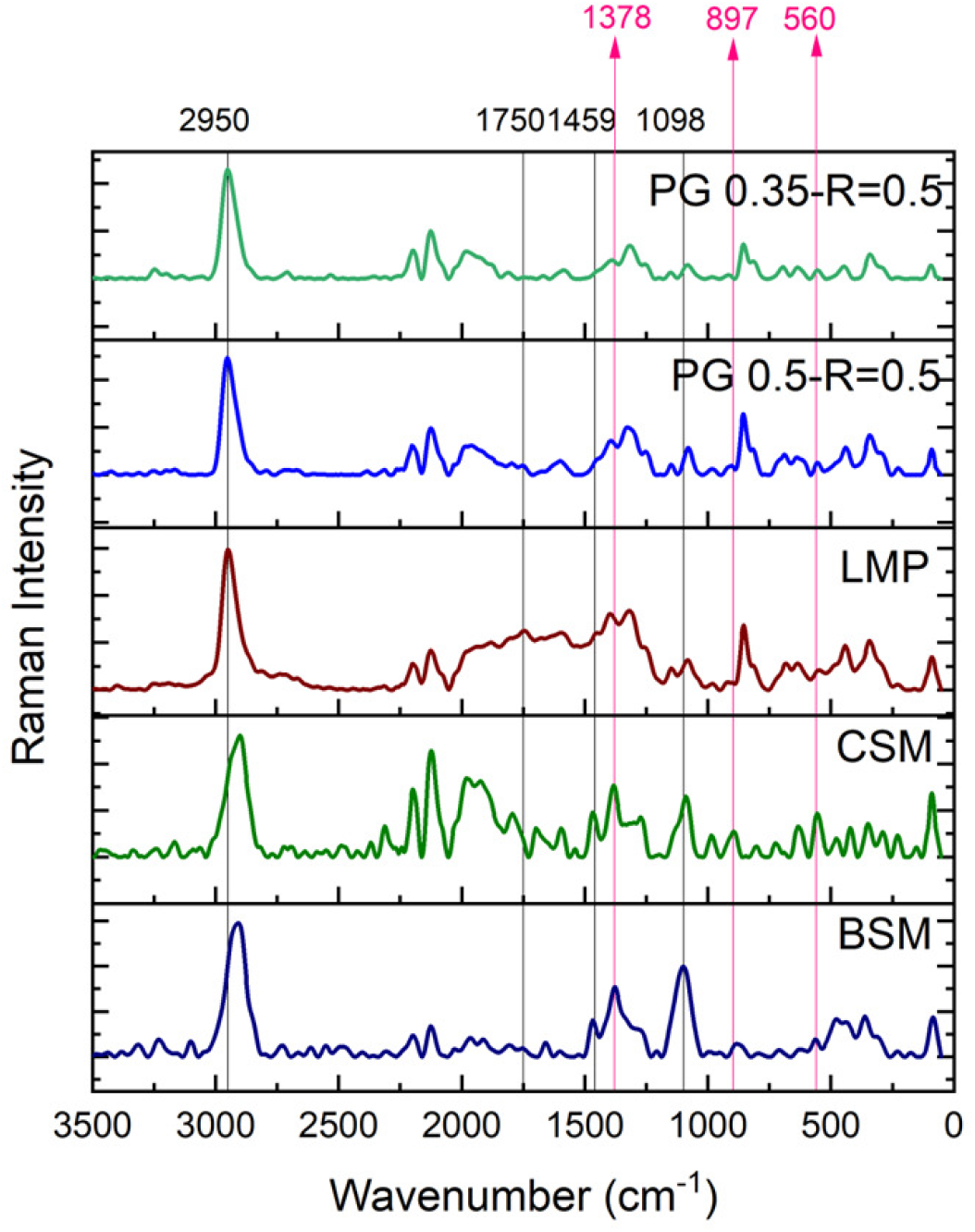
Fourier Transform Raman spectra of BSM, CSM, pectingels and LMP. The common peaks among pectin gels, pectin and seed mucilages (2950, 1750, 1459, 1098) are shown with black lines and the distinct peaks observed in seed mucilages (1378, 897, 560) are shown with coloured lines.

#### 3.2.2 Mechanical response

The results of LAOS rheology experiments on seed muciilges and calcium cross-linked pectin gels are shown in Figure 5. Among the different low methoxy pectin-Ca gels, the gels with 0.5 and 0.35 wt% pectin and R = 0.5 showed comparable zero shear storage modulus 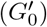 values to that of BSM and CSM respectively (Figure 5).

**Fig. 5.**
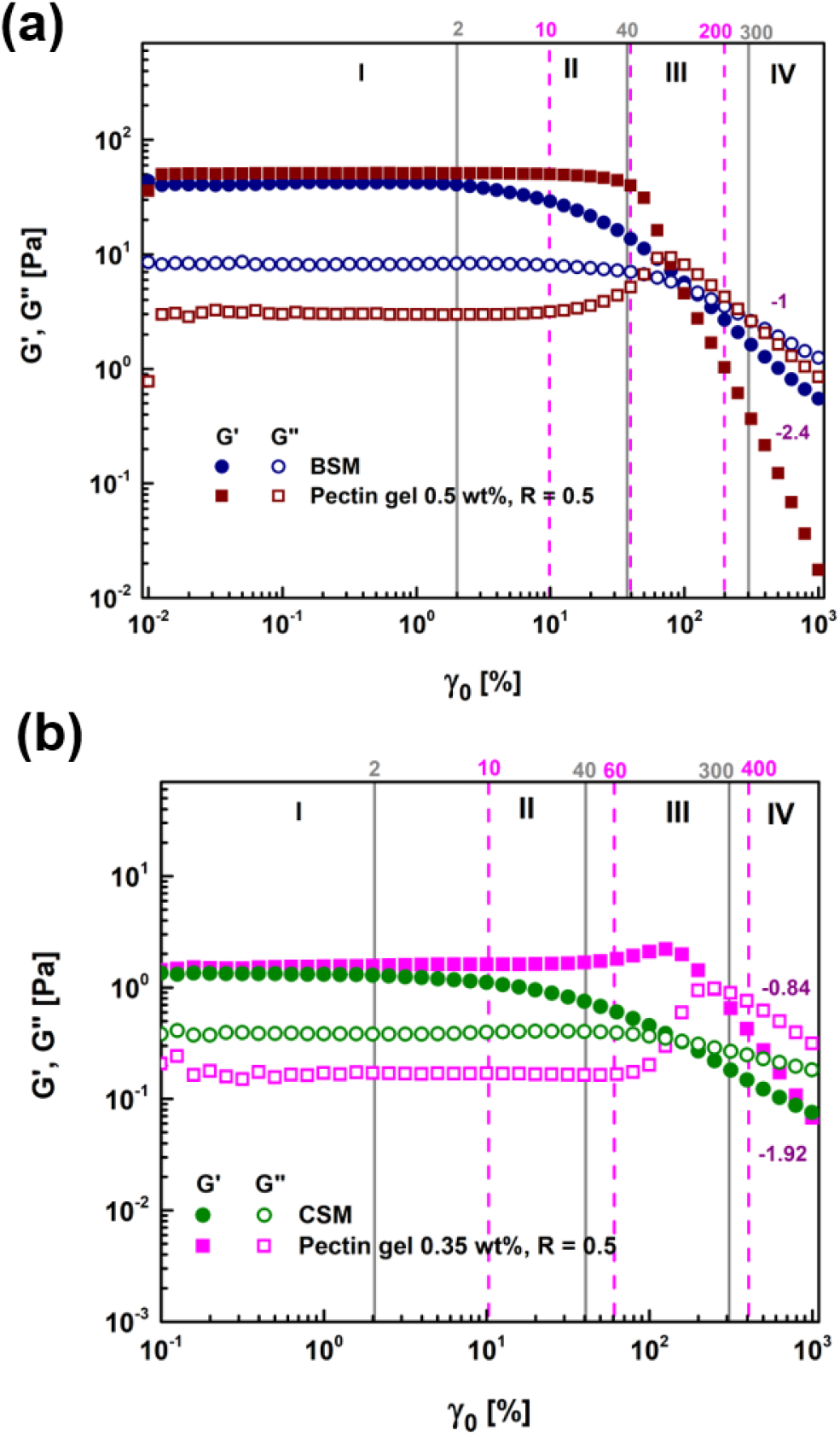
Comparative LAOS response of pectin gels along with seed mucilages (a) response of pectin gel of 0.5 wt% and R = 0.5 along with the response of BSM and (b) response of pectin gel of 0.35 wt% and R = 0.5 along with the response of CSM. The vertical lines represent different LAOS regions. The onset and extent of LAOS regions are different for seed mucilages (gray lines) and pectin gels (magenta dashed lines).

The linear viscoelastic limit for pectin gels is one order of magnitude higher than that of the seed mucilages. The overshoot observed in *G*^*′′*^ in region II is prominent in the case of pectin gels, weak in the case of CSM and absent in BSM. The limit of the linear viscoelastic region and the presence of overshoot are shown to be related to the presence of egg box bundles in the HG1 region of pectin chains in the presence of Ca [17]. It is interesting to observe that the non-linear region in seed mucilages occurs at much lower strain limits indicating possible differences in the microstructure of the mucilage or type of pectin and its crosslinking in the mucilage. In region IV, *G*^*′*^ ∝ (*γ*_0_)^*a*^ and *G*^*′′*^ ∝ (*γ*_0_)^*b*^. The values of *a* and *b* for pectin gels (Table 1) are higher compared to seed mucilages. Weak/no overshoot in *G*^*′′*^ and gradual decay of moduli at higher deformations in seed mucilages imply the differences involved in their composition and microstructure which might be the reason for the differences in observed response. However, at higher strain amplitudes, due to the presence of higher harmonics, *G*^*′*^ and *G*^*′′*^ data needs to be interpreted using other methods such as, intracycle (within a cycle of oscillation) stress-strain analysis using Lissajous-Bowditch (L-B) plots.

**Table 1.** Comparison of the mechanical response of seed mucilages and tack behaviour of swollen seeds with equivalent pectin gels. Values are obtained from Figures 5, 6, 7 and 8 and 9

| Property | BSM | CSM | PG 0.5 wt% | PG 0.35 wt% |
| --- | --- | --- | --- | --- |
| $G'_0$ [Pa] | 40 | 1.4 | 50 | 1.5 |
| Linear strain limit [%] | 2 | 2 | 10 | 10 |
| Overshoot in $G''$ | Absent | Minor | Significant | Significant |
| Yield strain [%] | 125 | 160 | 80 | 250 |
| a, b (decay coefficients) | -1.0, -0.63 | -0.8, -0.3 | -2.4, -1 | -1.92, -0.84 |
| Stiffening region [%] | 10-1000 | 30-1000 | 25-80 | 40-150 |
| Stiffening ratio in region IV | increases, $> 0$ | decreases, $> 0$ | $< 0$ | $< 0$ |
| Peak tack force [mN] | -87.96 | -27.77 | -55.26 | -26.78 |
| Debonding length [mm] | 9.16 | 8.94 | 7.06 | 6.51 |

### 3.3. Unique strain stiffening behaviour - a key feature for mechanical response of mucilage?

The L-B plots give better insights into the microstructural contributions of the mucilage from the observed LAOS behaviour. The L-B plots for BSM, CSM, and equivalent pectin gels with 0.5 wt% pectin and R=0.5 and 0.35 wt% pectin and R=0.5, at four strain amplitudes are shown in Figure 6. Various viscoelastic properties of the mucilages obtained from these plots are summarised in Table 1. The strain amplitudes represent the four regions of the LAOS shown in Figure 5.

**Fig. 6.**
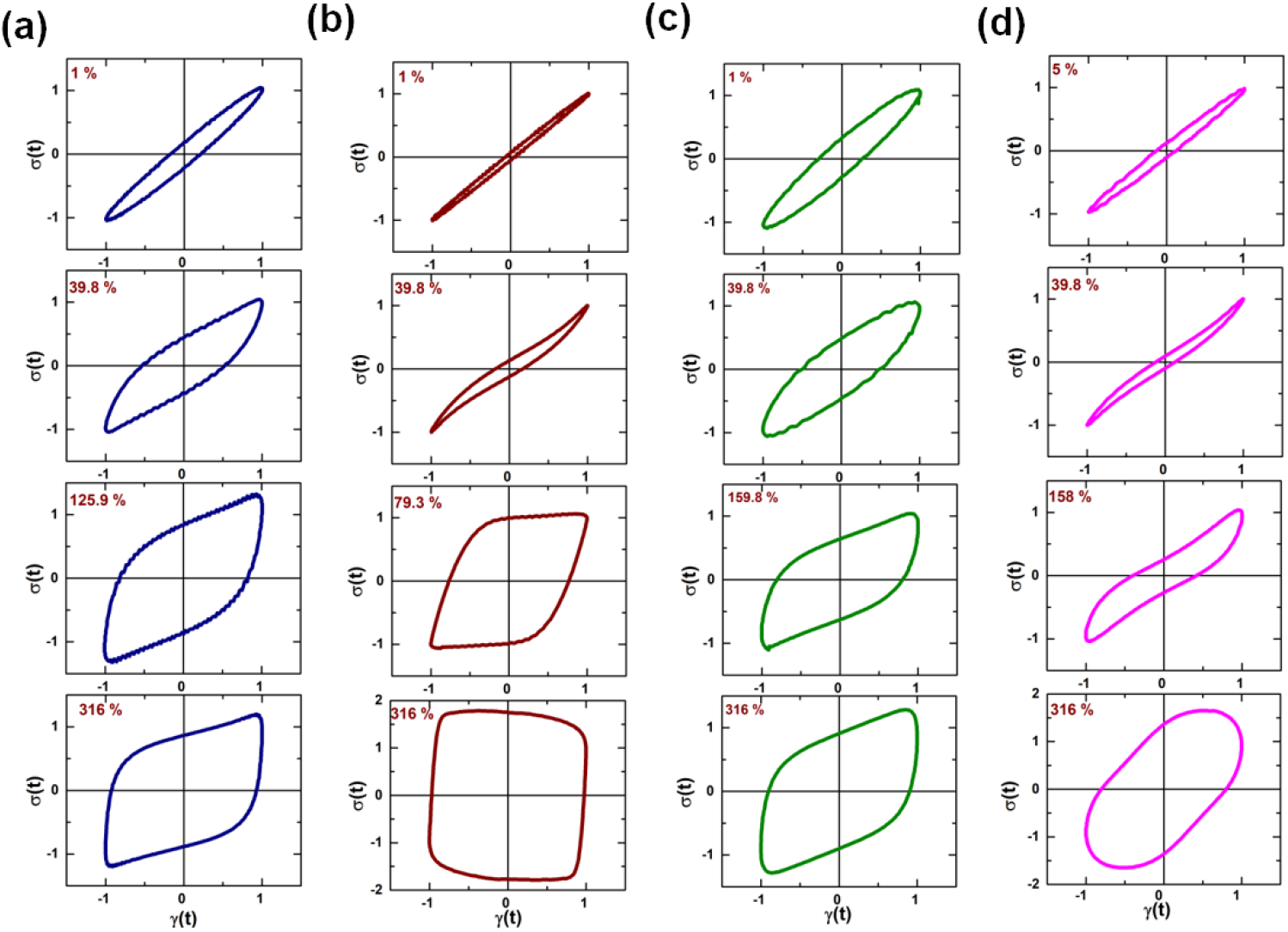
Lissajous plots for (a) Basil Seed Mucilage (BSM) (b) BSM Equivalent pectin gel 0.5 wt%, R=0.5 (c) Chia Seed Mucilage (CSM) and (d) CSM equivalent pectin gel 0.35 wt%, R=0.5. Strain regions are seletced based on Figure 5

The area under the L-B plots indicate the energey dissipation processes during the strain cycles. Both Basil and Chia seed mucilages show larger dissipation, especially at low shear in comparison to the pectin gels. This indicates the fundamental difference in the microstructure of the mucilages in comparison to the pectin based gels. In addition to this, the shape of these plots is elliptical within the linear viscoelastic region. However, beyond the linear limit, the L-B plots become non-elliptical indicating non-linear response of the material system. The shape and slope of the upward curve of the L-B plot (positive-half cycle) is indicative of another material behaviour, strain stiffening/strain softening. The strain stiffening/softening ratio, S for the mucilage and pectin gels (Equation 3) was calculated using MITlaos software [55] and is shown as a function of strain amplitude in Figure 7. 0*<*S*<*1 implies strain stiffening and S*<*0 implies strain softening in the material system. The S values and the strain stiffening/softening regions for pectin gels are comparable to that of the results reported previously [17]. The differences in the galacturonic acid content, degree of esterification and the source of low-methoxy pectin did not change the qualitative nature of the observed strain stiffening response in a significant way.

**Fig. 7.**
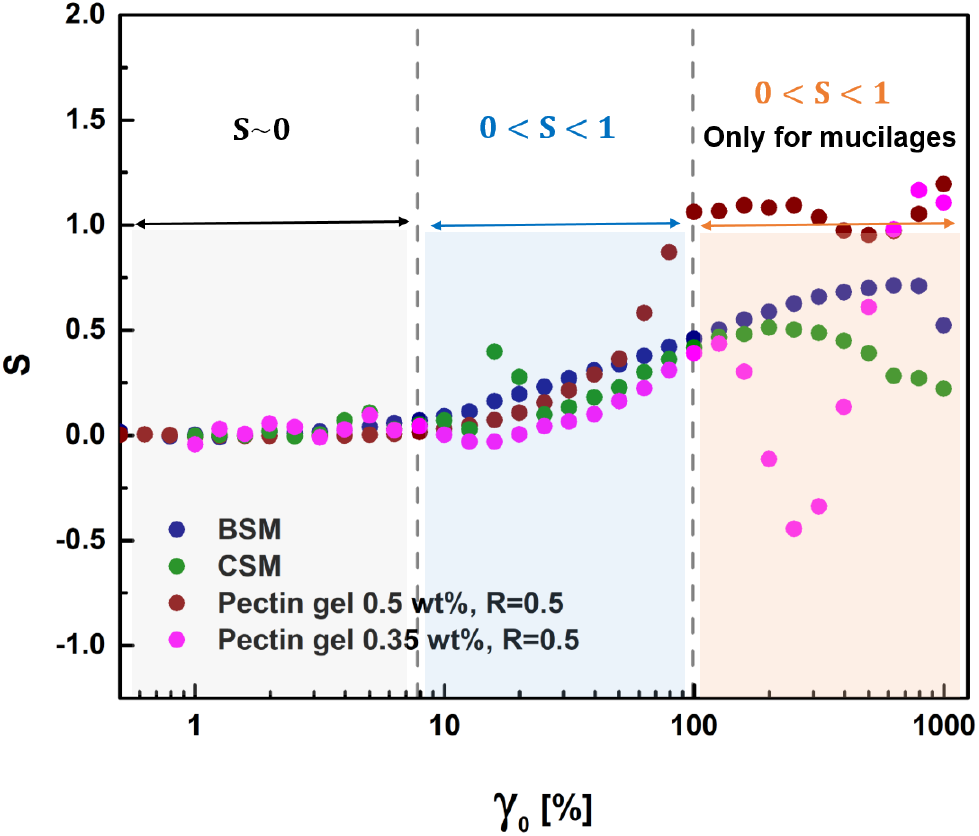
Strain stiffening ratio, S during large amplitude oscillatory shear for BSM, CSM and pectin gels. The dashed lines represent the different regions of strain stiffening of these gels.

The characteristic behaviour of both the mucilages and pectin gels, across their respective LAOS regions is interpreted using L-B plots. In region I (linear), the L-B curves are elliptic for all the systems (Figure 6) and S ∼ 0 (Figure 7) for both the mucilages and pectin gels implying linear response. In region II, for all the systems, L-B curves show concave upward nature and 0*<*S*<*1 indicates the strain stiffening behaviour. Even though the L-B curves appear steeper for pectin gels in this region, the S values for the seed mucilages are comparable with those for the pectin gels. At the end of region III and in region IV, in the case of pectin gels, the shape of the L-B curve changes and the S values are either *<*0 or *>*1. Interestingly, in the case of seed mucilages, the shape of L-B curve remains nearly the same with 0*<*S*<*1. It is important to note that S values should always be less than 1, i.e. S*>*1 has no physical meaning and cannot be interpreted as either stiffening or softening. This shows that beyond the cross-over strain amplitude, pectin gels do not show strain stiffening response, whereas, seed mucilages continue to exhibit strain stiffening even at larger strains. The area of the L-B plot indicates the energy dissipated in the material during deformation. It can be observed that for all the systems studied, the area of the L-B plots increases with increasing strain amplitude indicating increasing strain energy dissipation. In region IV (beyond the crossover), the energy dissipation is higher in the case of pectin gels. Similarly, the slope of the decay (*a*) of *G*^*′*^(*γ*_0_) in this region is also higher for pectin gels than seed mucilages as discussed earlier. This also points to the differences in the microstructure of the mucilage with that of the pectin gels and the processes contributing to the dissipation.

The transient Cole-Cole plots obtained from transient moduli are analysed using Sequence of Physical Processes (SPP) software [56] and are shown in Figure 8. The deltoid plot is given for three different strain amplitudes each corresponding to the non-linear regions II, III and IV of the LAOS. Changes in the position and the area of the deltoid with increase in strain amplitude are quite informative about the structure of the seed mucilages. At lower strain amplitudes (*γ*_0_ = 39.8% in Figure 8), major area of the deltoid lies below the 45° line, where 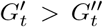 indicating more elastic nature of the gels. With increasing strain amplitude, the major portion of the deltoid shifts above the 45° line, where 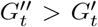 indicating a shift to fluid-like behaviour of the gels and also the yielding response. At higher strain amplitudes (region IV), the deltoid is completely located above the 45° line indicating more viscoelastic fluid- like response, especially in the case of pectin gels. However, in the case of seed mucilages, even at these higher amplitudes, some part of the deltoid still lies below the 45° line (in the shaded region) indicating the elasticity exhibited by the seed mucilages. This is also evident in the Lissajous curves, where the seed mucilages exhibit strain stiffening even at higher strain amplitudes, unlike the pectin gels. The significant observation from this analysis is that the seed mucilages have the capacity to maintain the elastic nature even at large strains.

**Fig. 8.**
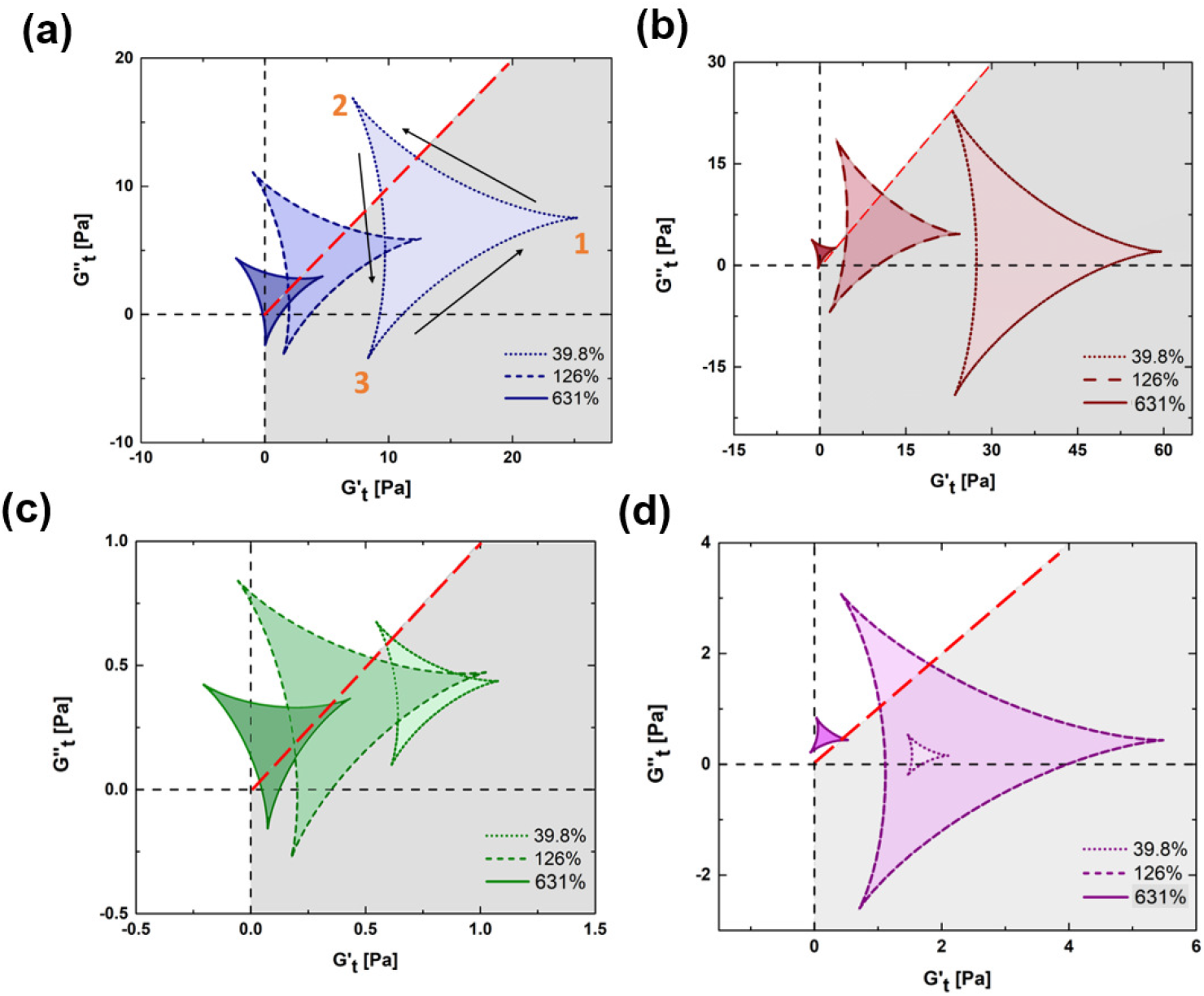
Transient Cole-Cole plots for (a) BSM (b)Pectin gel 0.5 wt%, R=0.5 (c) CSM and (d) pectin gel 0.35 wt% R=0.5. The black dashed lines represent the X and Y axes passing through origin. The red dashed 45° line represents flow condition, 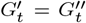. The shaded region represents 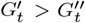 indicating elastic dominant region. The arrows indicate the path of deltoid and thus transient timescales such as yielding and recovery.

### 3.4. From wet tack and adhesion strength to seed anchroage?

Wet tack measurements carried out on the rheometer can give insights into the adhesive/debonding nature of soft materials. The wet tack response of swollen basil and chia seeds is shown in Figure 9(a) along with pectin gels of comparable gel properties. The normal force graph has two phases, an initial increase in negative normal force followed by a decrease in the normal force. The increase in negative normal force is due to the tension exerted by the material on the upper plate of the rheometer during retraction. This results in a peak force which is the maximum pull off force exerted by the material. After reaching the maximum pull off force, the normal force decreases as the material begins to detach from the plate. The area under the normal force and time curve (Figure 9(a)) gives the adhesion strength [Ns] of the material. Adhesion strength, peak tack force and detaching/debonding length of the swollen seeds and pectin gels are shown in Figure 9(b). Swollen basil seeds with mucilage have higher peak force and higher adhesion strength followed by equivalent pectin gel of 0.5 wt% R=0.5. Swollen chia seeds exhibit lower peak force and adhesion strength compared to basil seeds. Pectin gel of 0.35 wt% R=0.5 shows the lowest adhesion strength. However, during retraction of the upper platen, the swollen basil and chia seeds show higher detachment length than the pectin gels. This could be indicating that the cellulosic mucilages have higher wet tack strength and require stronger dislodgement forces to pull off from surfaces in comparison to pectin gels. The relation between the structure of the mucilage and its mechanical behaviour under shear are discussed further in detail.

**Fig. 9.**
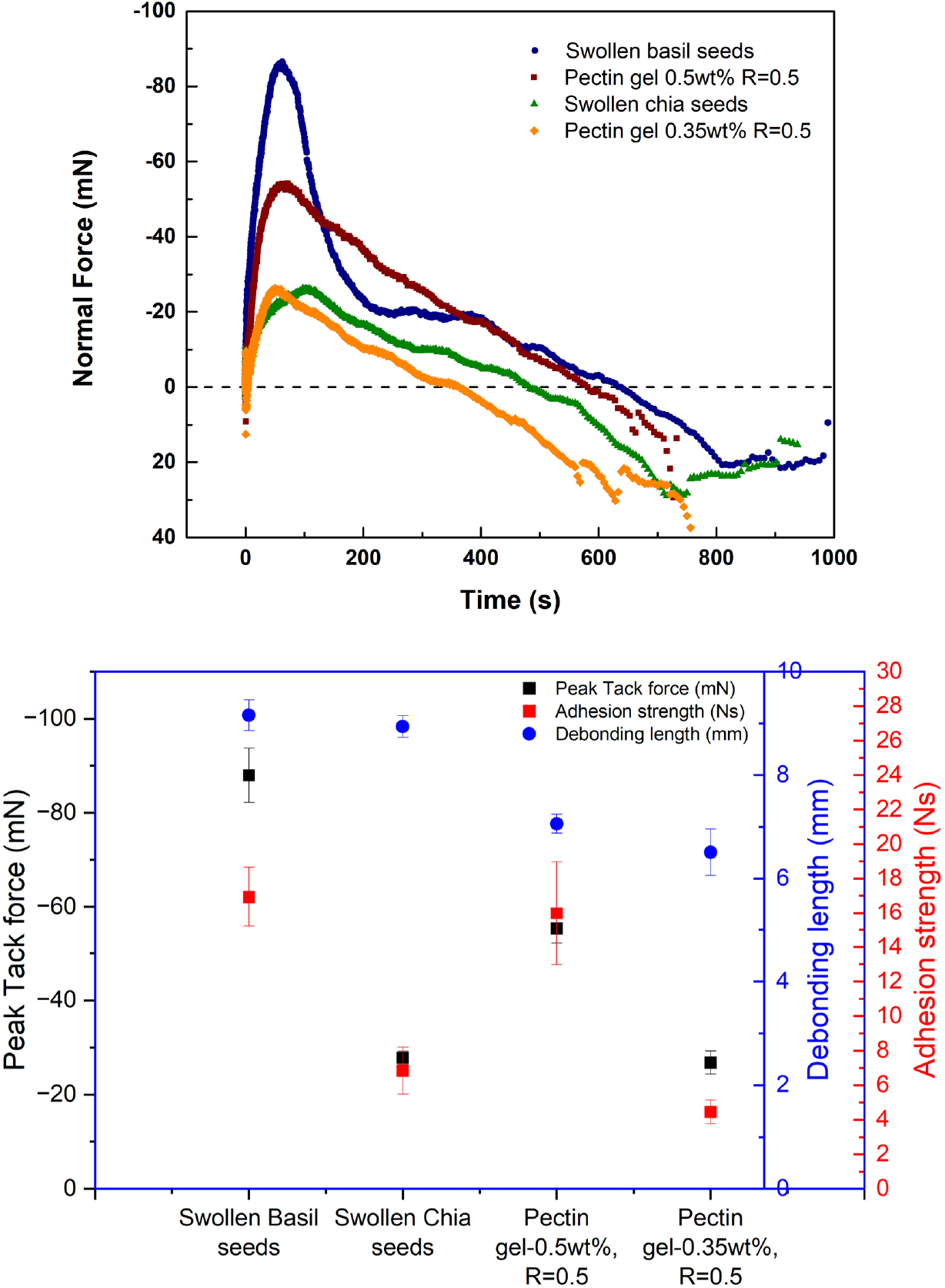
(a) Tack response of swollen basil seeds, chia seeds and pectin gels (b) Comparison of peak tack force, debonding length and adhesion strength

## 4. Discussion

### 4.1 Role of microstructure of mucilage in strain stiffening and wet tack behaviour

Since low methoxy pectin forms the matrix in mucilages, a comparative analysis of the rheological response of the basil and chia seed mucilages and the two pectin gels, are summarised below:(Table 1 and in Supplementary Information Table S1)

- Although two pectin gels with comparable zero shear modulus (*G*^*′*^_0_) to that of the seed mucilages were selected to reflect the mucilage behaviour, the linear strain limits for both the seed mucilages are found to be lower compared to that of the pectin gels.
- Lesser decay in the G^*′*^ values with increase in strain amplitude and the recovery of elasticity are observed in the case of seed mucilages even above the crossover strain amplitudes, unlike in the case of pectin gels (Figure 5).
- Pectin gels show strong overshoot in *G*^*′′*^ curves. Chia seed mucilage shows weak overshoot and basil seed mucilage shows no overshoot in G^*′′*^ values (Figures 3 and 5).
- In the case of seed mucilages, strain stiffening starts at lower strain amplitudes and persists till larger strain amplitudes (10-1000%). However, pectin gels exhibit strain stiffening only over a short range of strain amplitudes (25-150%) (Figure 7).
- Seed mucilages and pectins gels show comparable strain stiffening ratios (S) at low strain amplitudes. However, at strain amplitudes *>* 150%, seed mucilages continue to exhibit strain stiffening while the pectin gels show fluctuating S values indicating the differences in the microstructure and mechanisms contributing to the strain stiffening characteristics. Energy dissipation is also higher for the seed mucilages especially in region I and region II of the LAOS response (Figures 6 and 7).
- Swollen basil and chia seed mucilages exhibit higher tack force and adhesion strength than the equivalent pectin gels (Figure 9).

These differences in observed mechanical response could be arising from the microstructure of the mucilages. The role of microstructure in strain stiffening and the wet tack behaviour are discussed further to show how the behaviour helps in seed dispersal mechanisms in the case of cellulosic mucilage.

### 4.2 Role of cellulosic mucilage in anti-telechory

Although it is inferred that both the seed mucilages show unusual strain stiffening behaviour and good wet tack strength, it needs to be correlated to the functional traits of the seeds especially in dispersal. Here we explain how these mechanical responses can be interpreted to understand the anti-telechory exhibited by these seeds [8, 13]. By analysing the differences between the responses of the mucilages and the pectin gels, the mechanisms and processes involved in the strain stiffening behaviour of mucilages can be elucidated. Eventhough the 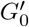 values of the mucilages and the corresponding pectin gels are comparable, the overshoot behaviour in *G*^*′′*^ values and strain stiffening characteristics are entirely different. Chia seed mucilage shows weak overshoot in *G*^*′′*^ and basil seed mucilage does not show any overshoot in *G*^*′′*^. This overshoot observed in *G*^*′′*^ in the case of pectin gels and alginate gels [17, 19] is attributed to the unzipping and reformation of the egg box-bundles formed by the macromolecular network of Ca crosslinked homogalacturonan chains present in the pectins. Hindrance to egg-box bundle formation has been shown to result in diminished overshoot or absence of overshoot in *G*^*′′*^ [17]. In the case of mucilages, the diminished overshoot or the absence of overshoot in *G*^*′′*^ could be originating from several factors, such as, the type of crosslinking (Ca or Mg), presence of monovalent cations that can hinder crosslinking by Ca ions (Na, K) etc. To verify this, EDS analysis of the mucilage was carried out and the results show the presence of divalent ions, such as, calcium (Ca^2+^) and magnesium (Mg^2+^) and monovalent ion, potassium (K^+^) in both basil and chia mucilages (Supplementary Information - Figure S1). The free uronic acids in the mucilage have the ability to crosslink in the presence of calcium and magnesium ions. However, pectin forms weak gels in the presence of Mg, compared to that of Ca [57]. The moduli (*G*^*′*^, *G*^*′′*^) of the mucilage indicate that they are preferably Ca-crosslinked gels (Supplementary Information - Figure S2). These suggest that diverse crosslinking mechanisms are plausible in mucilages when compared to Ca crosslinked pectin gels which could be leading to the observed differences in the rheological behaviour (LAOS-regions I and II), Figure 5).

The above descriptions are based on the microstructural features arising at the molecular scale, considering pectin based molecular networks alone. However, it is known that, the microstructure of the seed mucilages are constituted of hierarchical organization of different components [58]. The tubules found in the basil seed mucilage have thicker cellulosic fibrillary bundles which are connected by crosslinked pectin molecules and hemicellulose. Additionally, starch granules are also present along these cellulose fibrils(Figure 10). Chia seed mucilage has hierarchical structure with cellulosic fibrils connected by hemicellulose present in pectin gel matrix, but, lacks the starch granules and the thick cellulosic fibrillary bundles. Chia seed mucilage has no tubule formation either. In plant cell walls, it has been shown that pectin can bind with cellulose nanofibrils and alter the mechanical response [59, 60]. Hence, it is further examined to understand the role of the hierarchical structure of the mucilages in determining the mechanical response and the tack behaviour of the mucilage which may help in seed anchoring to the soil.

**Fig. 10.**
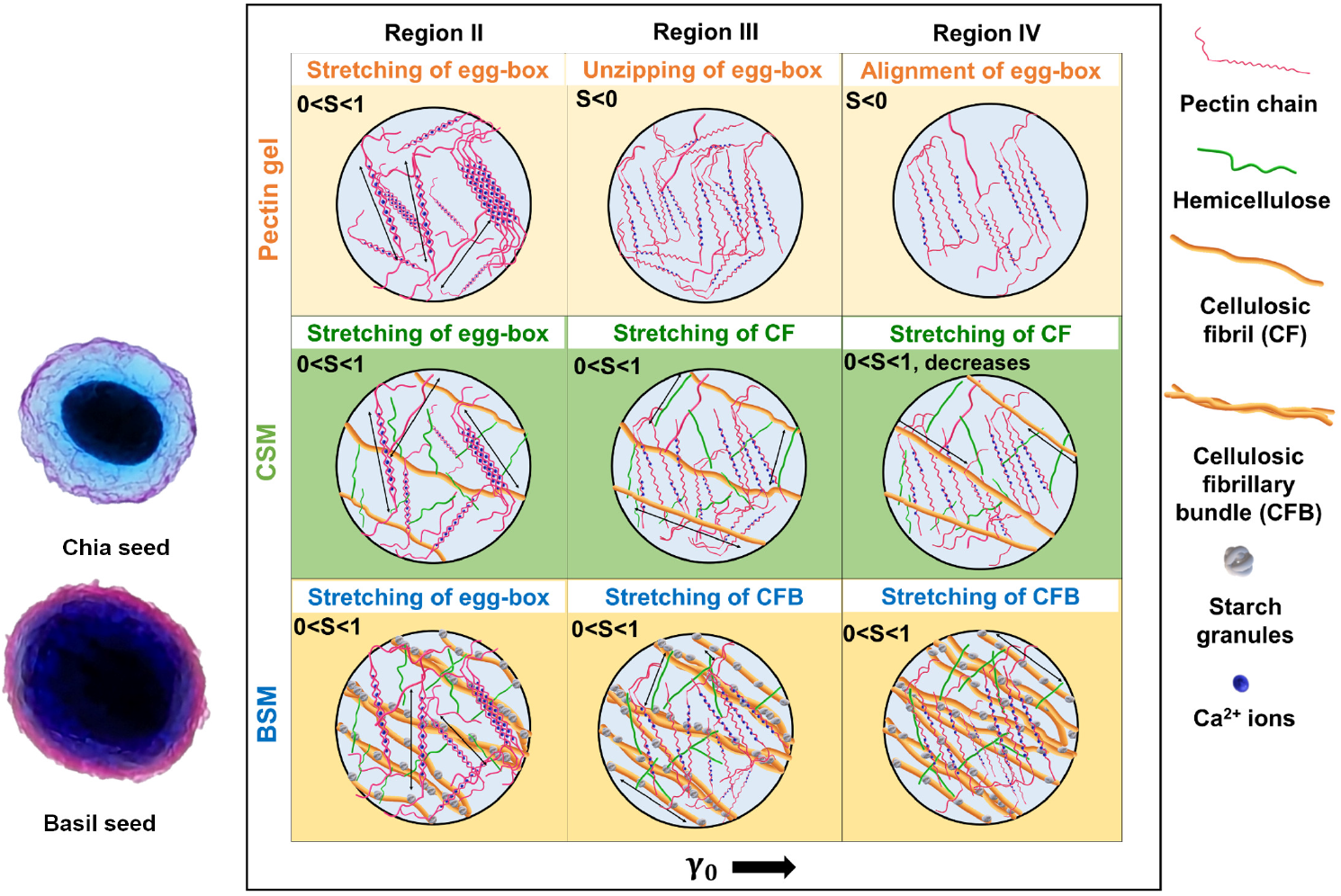
Schematic representation of the microstructural changes occurring in pectin gel and seed mucilages during deformation. The arrows indicate the stretching of egg-box bundles, cellulosic fibrils and their bundles. Swollen basil and chia seeds stained simultaneously using two dyes (Methylene blue and Ruthenium red) clearly shows higher amount of cellulose in basil seed mucilage in comparison to chia seed mucilage.

The unusual strain stiffening behaviour exhibited by two seed mucilages could originate from the microstructural features at multiple length scales present in the hierarchical structure. How the unique strain stiffening observed in the mucilages are different from the pectin gels is discussed below. In both the seed mucilages, the strain stiffening region is observed for a large strain range between 10 and 1000 % whereas, pectin gels show strain stiffening over a narrow range of 25 - 150% only. At lower strain amplitudes (*<* 100%), both the seed mucilages and the pectin gels show similar strain stiffening behaviour (Figure 7). This stiffening could be originating from the Ca- crosslinked pectin gel network present in mucilages and the pectin gels[17]. However, as the strain amplitude increases, only the seed mucilages show strain stiffening behaviour whereas, the pectin gels cease to show strain stiffening. The dissociation of pectin network through chain unzipping of egg box structures occurs and the role of pectin network in strain stiffening reduces. In fact, it has been also shown that at higher strain amplitudes, certain low methoxy pectin gels show strain softening behaviour [17]. However, observation of continued strain stiffening of mucilage highlights the fact that the stress transfer happens from pectin network to the hemicellulose and cellulose fibrils/fibrillary bundles under large strains. The possible strain stiffening mechanism in region III and IV could be originating from the stretching/bending of the cellulosic fibrils embedded in the pectin matrix. The difference in the strain stiffening response of basil and chia seed mucilages also clearly points to the different contributions arising from the cellulosic fibril/ bundles (basil) and cellulosic fibrils (chia). The schematic representation of the differences in the hierarchical structure and the microstructural changes during shear deformation of the seed mucilages and the pectin gels are shown in Figure 10. This emphasizes the fact that the hierarchical organization of components in seed mucilage and the systematic load transfer among different microstructural components helps the mucilage to withstand large shear deformations possible under water flow conditions. The mechanical response and adhesion behaviour of seed mucilages can be interpreted further in the light of seed anchorage or anti-telechory.

### 4.3 Mechanics of seed anchorage from rheological behaviour

In the natural environment, swollen seeds of basil and chia are subjected to various mechanical forces in the form of shear, compression, tension etc. These different external conditions could arise from events, such as the flow of water during the rains or watering the seeds after sowing. The typical shear stress experienced in these events by the seeds is in the range of 1.8-10.6 Pa and shear strains of 0.2-200% [11, 61]. These conditions can be simulated in rheological experiments using LAOS mode. The role of the extraordinary strain stiffening response observed in the case of basil and chia seed mucilages, in anchoring the seeds to the soil surface is further reinforced from the wet tack test results. The observed tack response also suggests that the mucilage layer helps anchor the seed strongly to the substrate and prevent its dislodgement by allowing large scale deformation of the mucilage without separating from the seed kernel. In the case of basil seed mucilage, tubule like structures are present which has long cellulosic fibril bundles along with hemicellulose and starch granules. Therefore, results from the wet tack experiments using whole swollen seeds indicate that the unique microstructure serve specific functions in nature. The tubules present in basil seed mucilage may help in additional anchorage by penetrating into the porous soil media. The stretching of the mucilage during shear events also enhances the area of contact with the soil. This response may even allow the mucilage to act like a cytoskeleton and protect the seed from rupture and dislodgement during large erosive forces. These inferences suggest that the microstructure and its hierarchical arrangement in cellulosic seed mucilage play crucial roles in seed physiology and seed dispersal.

## 5 Conclusions

Though the type of mucilage is known to affect the seed dispersal mechanism in myxodiaspory, in this work we show, the mechanism by which cellulosic seed mucilages help in seed anchoring and dispersal. When exposed to erosive forces such as those originating from the flow of water, cellulosic seed mucilage is found to prevent seed dislodgement by anchoring the seed to the soil/substrate. To understand the possible mechanisms by which mucilage supports seed anchorage, the mechanical responses of the two cellulosic seed mucilages - basil and chia, were analysed using both small and large amplitude oscillatory shear (SAOS and LAOS) rheology and wet tack studies. Similarities are observed in the qualitative mechanical response of both the mucilages exhibiting weak gel-like behaviour under SAOS and strain stiffening at large deformations. The response of seed mucilage to mechanical deformation is distinctly different from that of the model pectin-Ca gels studied. Especially, under large deformations, mucilage is capable of maintaining the elasticity and strain stiffening behaviour, unlike the pectin gels that undergo rupture and softening. Both the basil and chia seed mucilages show capability to recoil after larger deformations. This is also a unique feature of the seed mucilages when compared to the pectin gels. The presence of nano-scale structures such as cellulosic nanofibrils and their bundles interconneted by crosslinked pectin and hemicellulose in mucilage helps in withstanding large deformation through strain stiffening. The anchoring nature of the mucilage is established using wet tack experiments on a rheometer. The increased pull- off force exhibited by the mucilages during retraction is through strain stiffening of the cellulosic fibrils and this is unraveled using rheology. Strong seed anchorage which helps in anti-telechory is possible due to the contributions from the hierarchical structures present in seed mucilages resulting in more contact with the soil substrate. Our studies also suggest that rheology can be a novel approach to understand the physiology of plant based soft gels *in vitro*.

## Supplementary information

Supplementary Information

## Acknowledgements

This work is supported in part by funds from the Science and Research Engineering Board (SERB), India.

## Declarations

No declarations

