## Supplementary material for "Role of microstructure of cellulosic mucilage in seed anchorage: A mechanical interpretation": Supplimentary data

#### 1 Elemental analysis of seed mucilage

The Energy Dispersive Spectra (EDS) of both basil and chia seed mucilage (BSM and CSM) are shown in Figure 1. It confirms the presence of divalent ions such as calcium ($Ca^{2+}$) and magnesium ($Mg^{2+}$) and monovalent ions such as potassium ($K^{+}$) in both the seed mucilages. This indicates that the pectin present in the seed mucilages is possibly crosslinked through the calcium ions and forms the gel matrix in the seed mucilage.

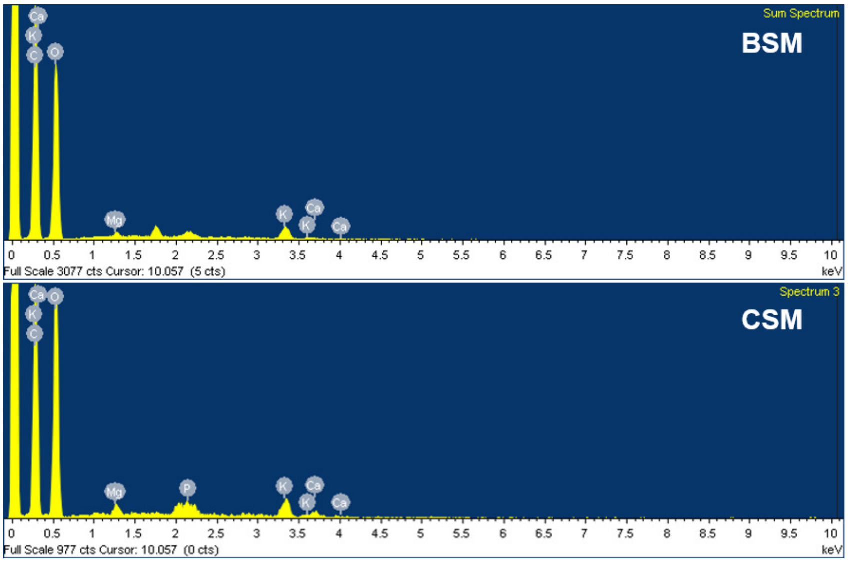


**Fig. 1** Elemental composition of (a) basil seed mucilage, BSM and (b) chia seed mucilage, CSM

#### 2 Seed mucilages - Ca or Mg crosslinked gels?

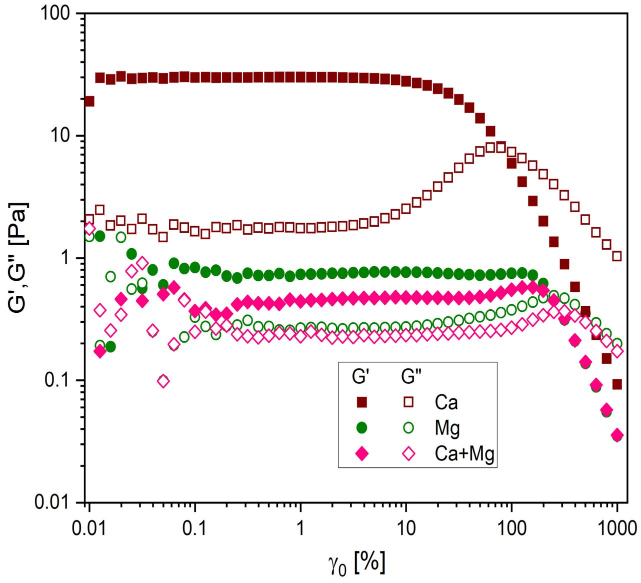


**Fig. 2** Large amplitude oscillatory shear (LAOS) response of pectin gels crosslinked by Ca, Mg and combination of both

Since both divalent ions (Ca and Mg) are present in the seed mucilage(Figure 1), the possible crosslinking ions is established from the response of the large amplitude oscillatory shear (LAOS) experiments on low-methoxy pectin crosslinked with Ca and Mg. The response for these three gels is shown in Figure 2. It can be observed that the zero shear modulus ($G'_0$) of pectin-Mg gel is lower than that of pectin-Ca gels. However, the seed mucilage has modulus values comparable with that of Ca crosslinked systems than Mg crosslinked systems. Additionally, the overshoot in $G''$ is not prominent in Mg crosslinked systems. The strain stiffening region of the pectin-Ca gels is 25-150% strain amplitude whereas, the stiffening region of pectin-Mg gel is in the range of strain amplitudes, 100-300%. It can be observed that pectin gels crosslinked by both Ca and Mg, do not exhibit strain stiffening at higher strain amplitudes like the seed mucilages. This indicates that other microstructural components such as, hemicellulose, starch grains, cellulose fibrils and their bundles present in the seed mucilages have significant role in the strain stiffening process observed at large deformations.

### 3 Selection of equivalent pectin gels to represent seed mucilage behaviour

To select the pectin gels to mimic the seed mucilage behaviour, the rheological response of different pectin gels with varying pectin concentration and cross-linker ratio were studied. Different rheological parameters such as, zero shear modulus ($G'_0$), linear strain amplitude limit and strain stiffening region are given in Table 1. From these compositions, PG 0.35 wt%, R=0.5 and PG 0.5 wt%, R=0.5 are selected to represent chia and basil seed mucilages respectively based on the obtained zero shear modulus ($G'_0$) values and also strain stiffening behaviour.

**Table 1** Large amplitude oscillatory shear response of pectin gels (PG) with different pectin content and calcium concentrations along with basil and chia seed mucilage

| Pectin gel compositions | $G'_0[Pa]$ | Linear strain limit [%] | Stiffening region[%] |
|---|---|---|---|
| PG 0.2 wt%, R=0.5 | 0.5 | 20 | 50-250 |
| PG 0.2 wt%, R=1 | 10 | 20 | 15-100 |
| PG 0.35 wt%, R=0.25 | 0.2 | 30 | 70-100 |
| PG 0.35 wt%, R=0.5 | 1.5 | 10 | 40-150 |
| PG 0.35 wt%, R=1 | 100 | 4 | Nill |
| PG 0.5 wt%, R=0.25 | 0.7 | 31 | 39.8-100 |
| PG 0.5 wt%, R=0.5 | 50 | 10 | 25-80 |
| PG 0.8 wt%, R=0.25 | 10 | 12 | 15-100 |
| PG 0.8 wt%, R=0.5 | 300 | 3 | Nill |
| PG 0.8 wt%, R=1 | 1000 | 1 | Nill |
| Chia seed mucilage | 1.4 | 2 | 30-1000 |
| Basil seed mucilage | 40 | 2 | 10-1000 |
